# Quantifying the Invasive Secondary Metabolome

**DOI:** 10.1101/2021.12.03.471125

**Authors:** Jamila Rowland-Chandler, Ewan Salter, Suresh Babu, Gitanjali Yadav

## Abstract

Invasive plants drive ecosystem degradation through developing aggressive phenotypes that can outcompete native flora. Several hypotheses explain this, like the Evolution of Increased Competitive Ability hypothesis and the Novel Weapons Hypothesis, but none have been proven conclusively. Changes in plant metabolites are critical to these hypotheses, but complete invasive secondary metabolomes have not been quantified. Here, statistical and unsupervised machine-learning approaches were used to analyse chemotype-to-phenotype relationships in invasive and non-invasive populations in species *Ageratum conyzoides, Lantana camara, Melaleuca quinquenervia* and *Psidium cattleainum* and on a family level analysing *Asteraceae, Myrtaceae* and *Verbenaceae*. Invasive metabolomes evolved according to the EICA and NWH, involving optimisation of aggressive strategies present in native populations and local adaptation.

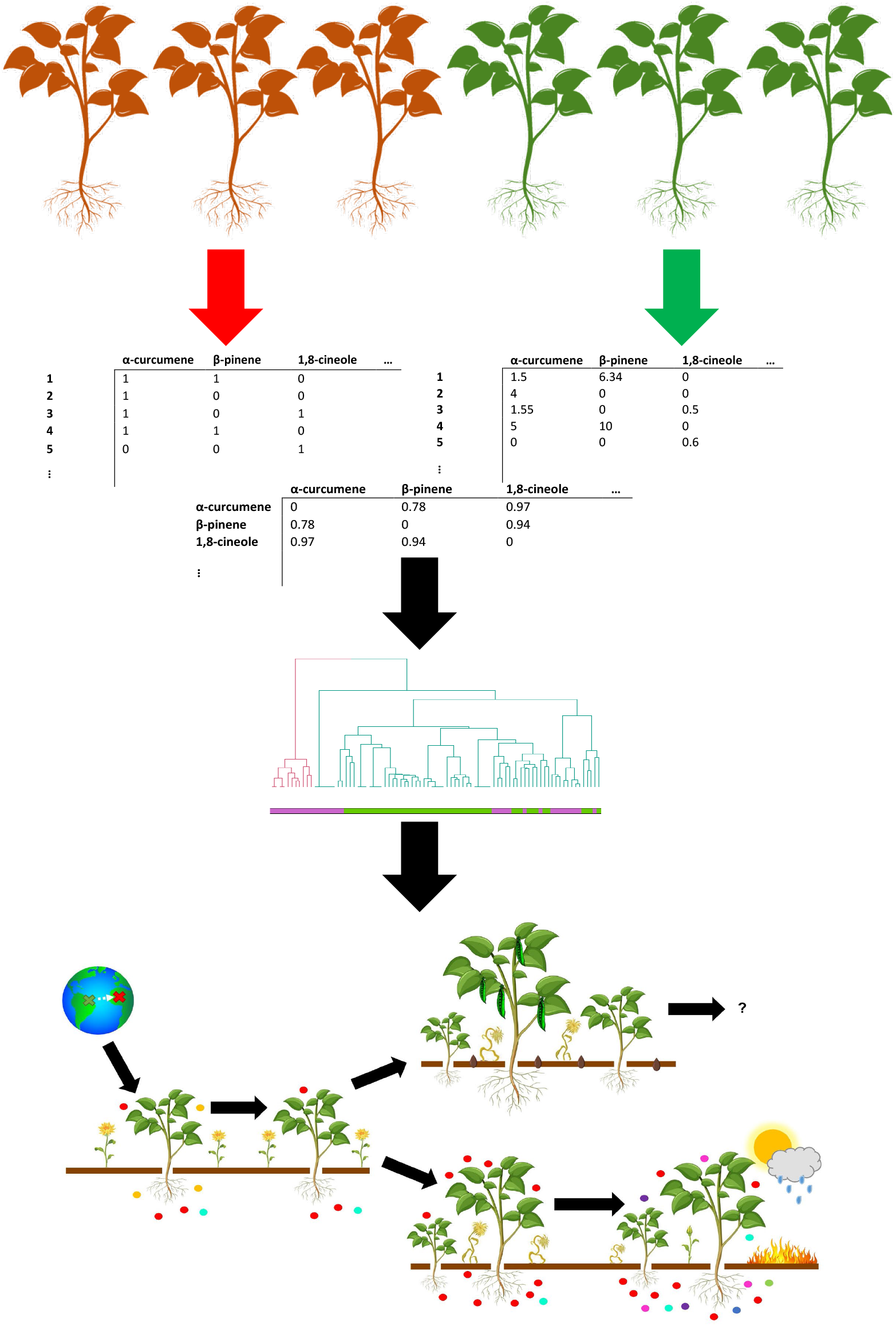

## 1. Introduction

Globalisation and anthropogenic movement have driven the mass emergence and spread of invasive plants. Invasives have higher fitness than native flora from experiencing less disease (Mitchel *et al*., 2003; Torchin *et al*., 2003), variably reduced herbivory and weakened competition from native plants. Therefore, invasives can proliferate wildly, so controlling these species is costly (Senator *et al*., 2017). Understanding invasive evolution in non-native plants is thus important in identifying which plants will become invasive and susceptible ecosystems that can be prioritised for protection by conservationists.

Two hypotheses addressing invasive plant evolution are the Evolution of Increased Competitive Ability hypothesis (EICA) and the Novel Weapons Hypothesis (NWH). EICA states plants in non-native environments will downregulate and lose defences against specialist herbivores and pathogens absent from non-native regions, allowing for resource reallocation to enhance growth and development to induce a rapidly proliferating invasive phenotype (Blossey and Notzold, 1995). NWH states some invasive plants produce novel compounds not found in native flora, involved in allelopathy (plant-plant “warfare”), generalist defence and other antagonistic strategies that increase competitive advantage (Cappuccino and Arnason, 2006). Thus, invasives outcompete native flora. Invasives will invest in producing these and new derived compounds to further increase competitive advantage (Callaway and Ridenour, 2004). Both hypotheses are underpinned by biochemistry. Plants secondary metabolomes have high chemical diversity, or chemodiversity, so plant-environment interactions, including those involved in invasive populations, are extensively mediated by secondary metabolites. In the EICA, divestment from specialist defence should involve divestment from related metabolite production and investment into different biochemistry. In the NWH, novel secondary metabolites mediate novel allelopathy and other plant-environment interactions in the invasive and these metabolites should increase in production and diversify. Therefore, metabolomes provide important insights into invasivity.

However, there is debate on whether the EICA or NWH are universally true across all invasives (Parker and Hay, 2005). This is in part because proving these hypotheses has been limited by lack of analysis of complete invasive secondary metabolomes. Previous research focused on assaying invasive chemical exudates on native flora and fauna, with varying agreement on whether the EICA or NWH are true (Lind and Parker, 2010; Siemann and Rogers, 2003), without identifying the compounds involved. Although some studies identified multiple important compounds (Cappuccino and Arnason, 2006), most focused on individual compounds or broad effects of chemical families only (Inderjit *et al*., 2006; Hull-Sanders *et al*., 2007). Therefore, important chemical variation of interest has been removed and combinatorial effects of multiple compounds have not been considered. Computational multi-omics approaches that can tackle large datasets could evaluate chemotype-to-phenotype relationships from invasive secondary metabolomes, so interdisciplinary approaches are needed to tackle questions in invasive plant evolution. Furthermore, very little research has studied the EICA and NWH together, which likely interact in many species (Qin *et al*., 2013; Zheng *et al*., 2015). Outside these major hypotheses, there has been limited study on long-term invasive secondary metabolism evolution after the emergence of aggressive phenotypes, and thus how competitive advantage may shift (Strayer *et al*., 2006). All these gaps in knowledge need to be addressed.

In this study, we set out to evaluate whether EICA and NWH hypotheses are true, either in conjunction or independently, through analysing secondary metabolomes, as these hypotheses are likely to be at least partially evident from chemistry alone. The native ecology of invasives may also determine to what extent strategies predicted by the EICA and NWH are implemented in invasive populations. Invasive secondary metabolome evolution should associate with environmental and ecological factors in non-native regions, explaining how invasives can proliferate in the long-term and why some ecosystems are more susceptible to invasive takeover. To investigate these hypotheses, a meta-analysis of the chemical composition of essential oils of invasive and non-invasive populations across several species was conducted, allowing the chemical evolution of the invasive from native populations to be studied. Unsupervised Machine Learning methods like cluster analysis were used to compare chemical composition between plants. This was appropriate for identifying underlying patterns in variation of compound diversity, production levels and chemical relationships between metabolites from large datasets. Similarity-based measures and robust statistical testing were also used to analyse trends and to tolerate the high variance present in datasets. From this analysis, we determined chemistry of all invasives studied followed both or individually the EICA and NWH to varying degrees, where invasives evolved to optimise pre-existing aggressive strategies whilst responding to some selection pressures in the non-native environment.

## 2. Methods

Data extraction and statistical and computational analysis were performed in the statistical computing software “R” (Version 4.0.2, http://www.r-project.org).

### 2.1 Data Collection

#### 2.1.1 Plant Chemical Profiles

Invasive plant species were identified through GISD “100 most invasive species” and EssoilDB (The Global Invasive Species Database; Kumari *et al*., 2014). *Lantana camara, Melaleuca quinquenervia, Psidium cattleainum* and *Ageratum conyzoides* were selected for data availability reasons. Essential oil chemical composition, or profiles, were extracted from EssoilDB source articles using web-trawling methods with packages XML, rvest and stringr or from raw datafiles (CABI 2020). Further profiles were collected manually from primary literature (Riaz *et al*., 1995; Philippe *et al*., 2002; Zoghbi *et al*., 2007; Monti *et al*., 2009; Nitin *et al*., 2010; Tesch *et al*. 2011; Castro *et al*., 2015; Kouame *et al*., 2018). Profiles with reasonably complete-looking metabolomes were selected so that the data partly reflected the complete population chemodiversity. Most of the data for plant families was extracted separately from EssoilDB and the compiled with the species data.

Compounds present and % amount of each in the essential oils (as determined by GC-MS), location and date of sampling where possible were extracted. Whether the plant was native, non-native or reported as invasive in the country of sampling were identified through GISD and CABI (The Global Invasive Species Database, CABI 2020).

#### 2.1.2 Environmental and Ecological Factors

Climatic data was collected from Wikipedia and forecast websites **(https://www.weatherandclimate.com, https://www.weather2visit.com, https://weatherspark.com)** from the closest major city to the plant sampling site, as data for rural areas was sparse. Average annual precipitation (mm), average monthly high temperature (°C), average monthly low temperature (°C), average monthly relative humidity (%) and elevation (m) were sampled. Flood risk was estimated from news articles and flood warnings. Disturbance data was obtained from global forest watch databases from the province the plant sampling site was located (Global Forest Watch): % total tree cover loss (2001-2019), total VIIRS alerts per annum (2020) and presence of anthropogenic activity.

Packages rgbif and Countrycode were used to measure *Plantae* and *Animalia* (for plant species data only) Species richness (SR) per country from which plants were sampled. This was calculated from number of unique species from each kingdom where occurrence coordinates were known in GBIF databases (GBIF.org). Per country data, +/-10yrs from plant sampling date where known, or from 1980-2020 where sampling date was not, was selected as occurrence records were too inconsistent to calculate SR of the year of plant sampling. This assumed most easily surveyable species in a country would be identified in a 20yr survey period, so the 20yr and 40yr records would be similar.

### 2.2 Data Normalisation

Species names were normalised using Taxise-package. Plant part sampled, invasivity and native/non-native status, flood risk and presence of anthropogenic activity were normalised to binary data through non-package associated code so that categorical data was consistently named. Country was converted to factorial data. Other data was kept raw and numeric. Compounds common names, given in the literature, were converted to Canonical SMILEs using packages webchem, rJava and rcdk. SMILEs were converted to 1024-bit Morgan Circular Fingerprints using packages rcdk and rcdklibs. Profiles and corresponding plant features, environmental and ecological factors were stored as feature vectors. One data matrix with binary presence and absence of compounds, the other with production rates as % amounts were constructed.

### 2.3 Computational and Statistical Analysis

Two matrices were produced – one for data by species, the other by family. The species matrix had information on compound production, whereas the family matrix was binary presence/absence data. Computation and statistical analysis were conducted on invasive and non-invasive populations for all species families except *A. conyzoides* and *Asteraceae*, where native and non-native populations were compared, as this species was recorded to be non-invasive in many countries in its non-native range. Methods applied to quoted invasive/non-invasive and non-native/native population sets were consistent.

#### 2.3.1 Initial Plots and Statistics

Individual chemical profiles were plotted using pheatmap-package. To measure the variation between all chemical profiles, thus determining if there was divergence between profiles, pairwise Bray-Curtis dissimilarity between profiles was calculated from binary presence/absence compound data using the vegan-package, then converted to a distance matrix and mean dissimilarity was calculated.

#### 2.3.2 Profile Size Analysis

Chemical profile size was calculated from total number of distinct SMILES per profile. Due to unequal variance, Kruskal-Wallis and Mann-Whitney non-parametric tests were used to compare profile size between species and invasive and non-invasive populations within and between species.

#### 2.3.3 Profile Clustering

Clustering algorithms were applied to presence/absence and % amounts profile datasets to assess divergence between invasive and non-invasive populations. Hierarchical clustering was performed per species using the factoextra-package with a defined cluster number: k = 2, using Euclidean distances and the Ward clustering algorithm. The clusters obtained were validated with the expected clustering – invasive and non-invasive – using the fpc-package to assess strength of similarity. Adjusted Rand Index was the similarity measure used. Hierarchical clustering was also used to assess divergence in chemical structural diversity between invasive and non-invasive populations. From a Tanimoto’s distance matrix of all compounds identified, pairwise distance between group centroids of profiles per species was calculated using ANOVA-like tests with usedist-package. Each group was the compounds present in a profile. A new distance matrix between profiles was constructed and clustered using factoextra-package with same methods as previously stated. The clusters obtained were validated with expected invasive/non-invasive clustering with methods pre-stated.

Annotated dendrograms were plotted with dendextend-package.

#### 2.3.4 Changes in structurally related compound production unique-to-population synthesis

To determine whether compound production differed between invasive and non-invasive populations from the species matrix, Kruskal-Wallis non-parametric tests were calculated for each compound shared between some invasive and non-invasive populations; profiles that did not produce the compound were denoted 0% production. Whether there was any chemical similarity between the compounds with altered production was also assessed to identify functional convergence within invasive and non-invasive profiles. K-medoids clustering was implemented on a Tanimoto’s distance matrix of all compounds to determine structural grouping between compounds using cluster-package. The optimal cluster number was found using the average silhouette method with the Euclidian distance metric using cluster-package. Exact tests were implemented to evaluate difference in proportion of compounds belonging to each cluster upregulated and downregulated in invasive compared to non-invasive populations. Chi-squared and exact tests were used to compare proportion of compounds per cluster unique to invasive and unique to non-invasive populations. For the family matrix, because the difference in sample sizes between populations was large, where possible a bootstrapping method was also used to compare equal sample sizes. Range and Average number of unique-to-population compounds and the p-values from multiple statistical tests were reported. Bootstrapping was iterated 10 times.

#### 2.3.5 Evaluating the impact of environmental and ecological factors on invasive chemical profiles

To assess whether there was invasive evolution in response to environmental and ecological factors in non-native regions, linear and generalised linear models (GLM) were run. Data from invasive populations of *L. camara* only, as no other species had sufficient data, on average compound production per cluster per profile was modelled using environmental and ecological factors in a robust linear model (RLM) using Mass-package. This was to determine whether chemical similarity and production levels associated with the local environment. The significance of explanatory variables was evaluated using Ward tests from sfsmisc-package. *L. camara* only and cluster 1 and 2 had sufficient data to be analysed. For the family data, compound enrichment per cluster for all families was modelled using a Poisson GLM to evaluate the effects of environmental and ecological factors. One model per cluster was created and within clusters models each containing one of three were created to ensure meaningful data was not removed from the analysis. All clusters were analysed bar cluster 6 due to 0 inflation.

## 3. Results

### 3.1 There is high variance in metabolome composition within species

The family and species analysis can be distinguished as follows: the species data contains data from A. conyzoides, L. camara, M. quinquenervia and P. cattleainum; the family data includes *Asteraceae* (*A. conyzoides* and other non-native and native species), *Verbenaceae* (*L. camara* only) and *Myrtaceae* (*M. quinquenervia* and *P. cattleainum*, plus some non-invasive species). Across all species and families there was high variation in the composition, chemical diversity and production levels of the chemical profiles collected (Fig. 1). Therefore, there is evidence of metabolomic divergence within species and families, which could be attributed to differences in evolution between invasive (or non-native) and non-invasive (or native populations).

**Fig. 1.**
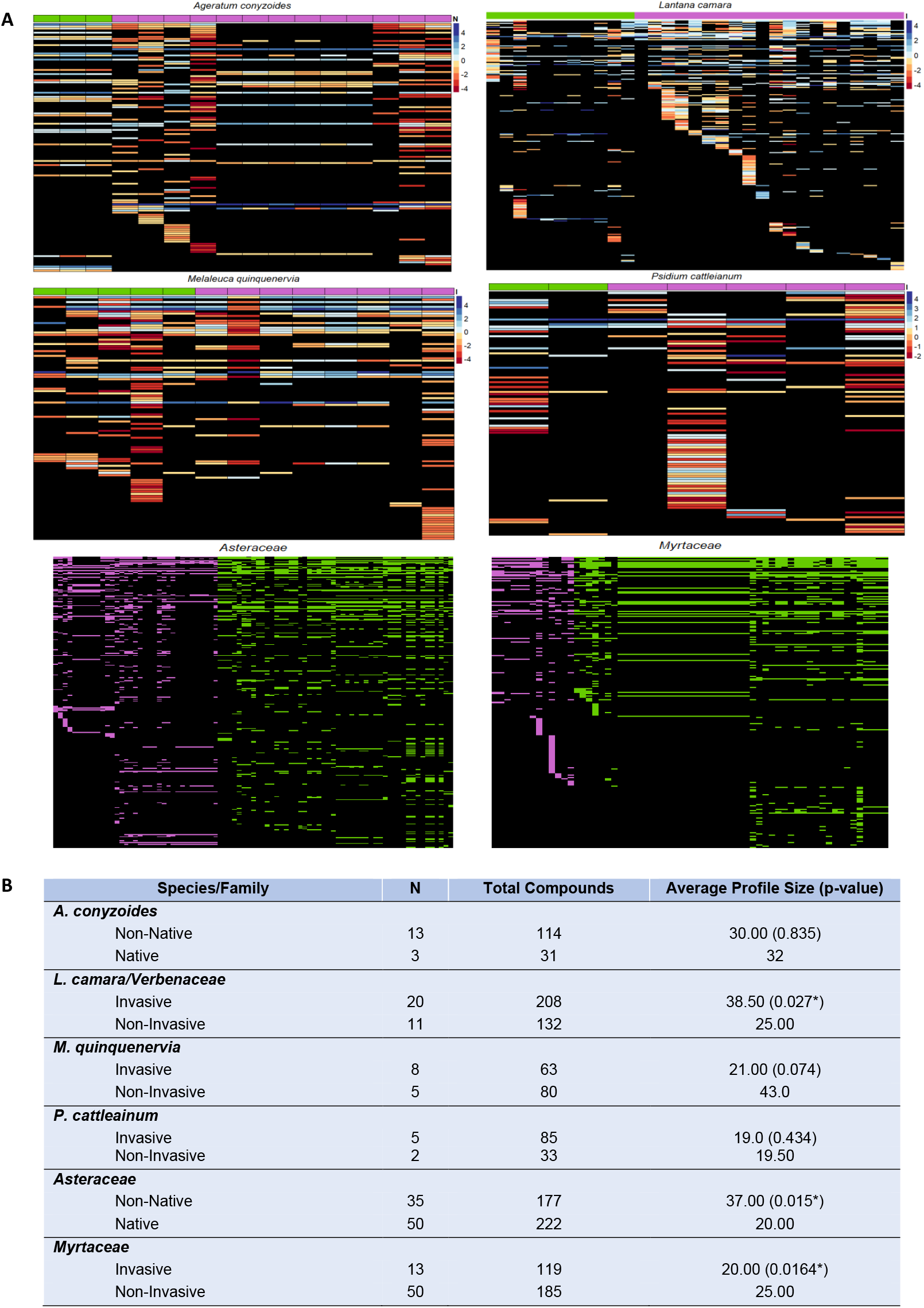
Summary of the chemical diversity observed in the chemical profiles. (a) Individual chemical profiles. Top and centre plot cell colours illustrate log(compound production) and the annotation bar shows which profiles are invasive or non-native (purple) and which are non-invasive or (green). For the bottom plots, coloured cells indicate compound presence and are coloured according to whether the profile is from an invasive/non-native (purple) or non-invasive /native populations. (b) Initial statistics from the chemical profiles between invasive or non-invasive or non-native and native populations. Average profile size is estimated from the median number of compounds per profile and p-values are from Kruskal-Wallis tests comparing profile size.

### 3.2 Broad chemodiversity-related strategies differ between species, but show evolutionary divergence between populations

According to the EICA and NWH, invasive chemical profiles may change in size and chemodiversity as compounds are lost and metabolic diversity may radiate during evolution in non-native environments. Therefore, convergent evolution in invasive populations can be initially assessed through investigating profile size and chemodiversity. Total chemodiversity was generally higher in invasive than non-invasive species profiles, but *M. quinquenervia* and families *Asteraceae* and *Myrtaceae* showed the opposite (Fig. 1b). There were no obvious trends in number of compounds produced per plant between invasive/non-native and non-invasive/native populations across all species (Fig. 1b) and there was no significant difference in profile size between populations when comparing all species and families (Kruskal-Wallis, H(1) = 1.10, p = 0.294; H(1) = 0.000640, p = 0.980). However, there were differences in profile size within taxa; *L. camara* and *Asteraceae* had expanded whereas *M. quinquenervia* and *Myrtaceae* reduced in invasive populations, all statistically significant (Fig. 1b). *P. cattleainum and A. conyzoides* had similar profile sizes between populations.

Therefore, invasives have diverged from non-invasive metabolomes, but evolutionary strategies are inconsistent across taxa

### 3.3 Invasive populations show chemical divergence, but share similarities in production levels and chemical properties of the secondary metabolome

2-Cluster analysis showed there was not strong segregation between invasive or non-native and non-invasive or native populations of all taxa when clustering profiles according to chemical composition chemical similarity and compound production (the last for species only). (Fig. 2a). On the other hand, from inspection, within clusters invasive and non-invasive profiles tend to cluster together, suggesting there is some consistency in profiles within populations. However, similarity measures did not show these trends, although from taxa with larger more robust datasets, like *L. camara* and *Asteraceae*, chemical similarity and production levels are stronger signals of invasivity than chemical composition, showing greater agreement with expected invasive/non-invasive clustering (Fig. 2b). The same was shown for production levels *M. quinquenervia* once biasing chemotype compounds were removed, and less obviously in *A. conyzoides* as clustering completely changed from chemical composition and similarity dendrograms, which were biased by the sampling of same compounds. Contrastingly, these trends in relation to chemical similarity and production were not seen in *P. cattleainum* and *Myrtaceae*. Therefore, generally metabolomes of non-native or invasive populations could show functional similarity.

**Fig. 2.**
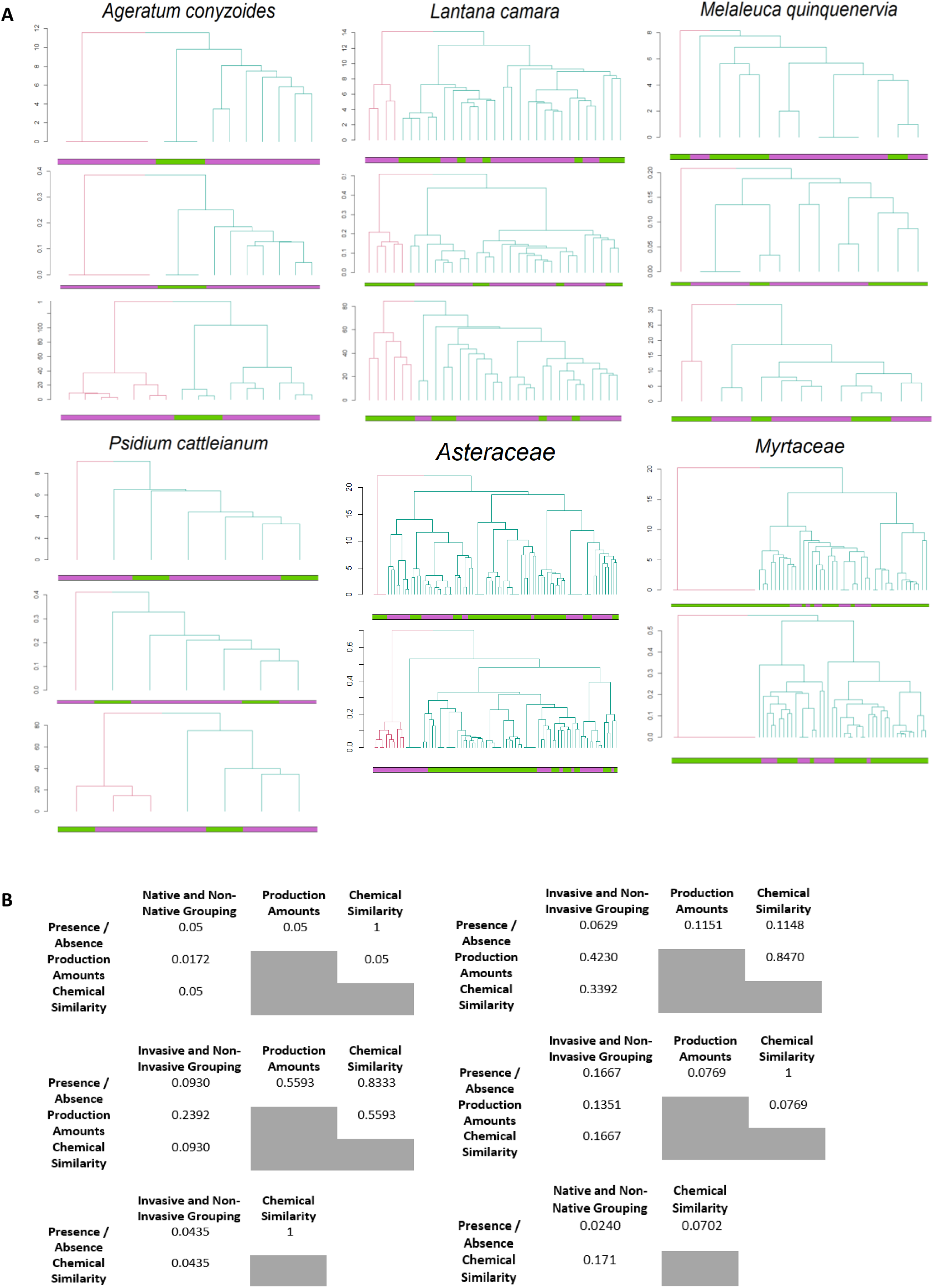
Clustering of invasive and non-invasive or non-native and native chemical profiles. (a) Hierarchical clustering analysis of chemical profiles based on presence and absence data (top), structural diversity of compounds (middle) and compound production level (%) data (bottom) in each profile (bottom). Profiles were clustered into two groups, shown by branch colouring, using Euclidean distance measures and the Ward Clustering algorithm and annotated to show which profiles were invasive/non-native (purple) or non-invasive/native (green). (b) Similarity matrices between true and expected invasive/non-invasive or non-native/native clustering and between the clustering for each data type. From top left, clockwise: *A. conyzoides* (*Asteraceae*), *L. camara* (*Verbenaceae*), *P. cattleainum, Compositae, Myrtaceae* and *M. quinquenervia*. Similarity between two-group clustering is calculated using a modulus adjusted Rand index.

Additionally, native/non-invasive populations may have undergone more similar chemical evolution than non-native/invasive populations. Native/non-invasive populations tend to segregate in larger sub-clusters than non-native/invasive populations and non-native/invasive profiles formed multiple small clusters often grouped with individual native/non-invasive profiles (Fig. 2a). This suggests individual invasive profiles show greater and more rapid divergent evolution. Alternatively, some native populations could have an “invasive-like” phenotype, so could be primed for invasivity once introduced to non-native environments. Thus, invasive populations may have metabolically radiated from each other, but show important similarities between compound production levels and the chemical similarity of profiles.

### 3.4 Production levels and unique-to-population synthesis of structural similar compounds reveals strong distinction between populations

Many compounds had different production rates between non-invasive/native and invasive/non-native populations in all species (Fig. 3b, 3d). A number of individual compounds had significantly altered production between populations in *A. conyzoides* and *L. camara*, but for many the difference was non-significant, and no compounds had significantly different production in *M. quinquenervia* and *P. cattleainum* (Fig. 4d). Furthermore, species differed in terms of the number of compounds upregulated and downregulated in invasive populations – *L. camara* and *M. quinquenervia* had more upregulated compounds, *A. conyzoides* and *P. cattleainum* had more downregulated. However, when compounds were clustered according to structural similarity, trends between species were observed. All species but *P. cattleainum* had an overrepresentation of cluster 1, but this was only significant for *L. camara* and downregulated compounds in *A. conyzoides* (Fig. 3b). Additionally, the compounds with significantly different production were mainly found in cluster 1. Deviation from expected cluster proportions was not shown for any other groups except cluster 3 in *L. camara*, which were significantly underrepresented. Therefore, invasivity could be driven by altering production levels of chemically similar compounds and by the additive effects of many rather than a few key compounds.

**Fig. 3.**
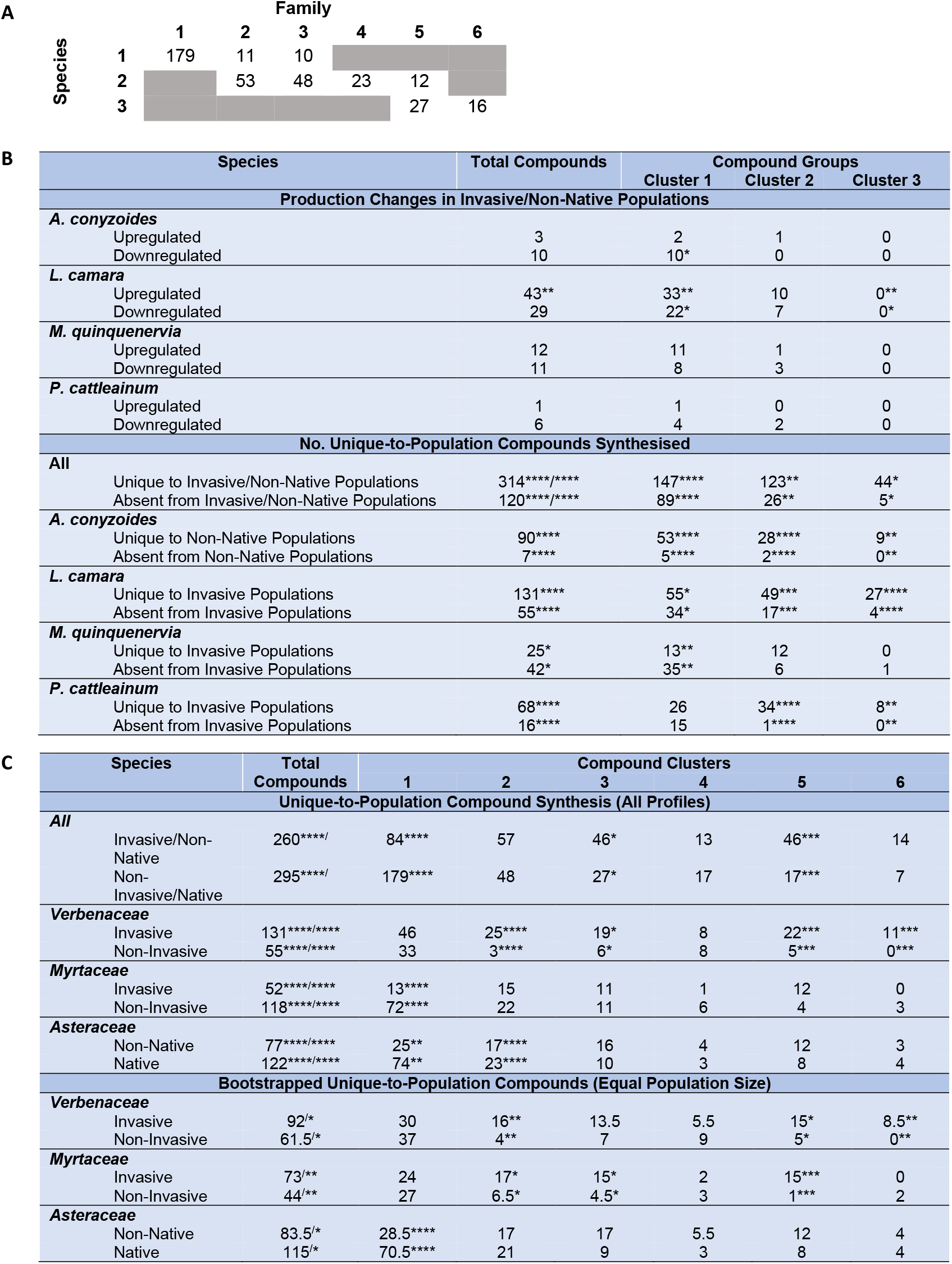
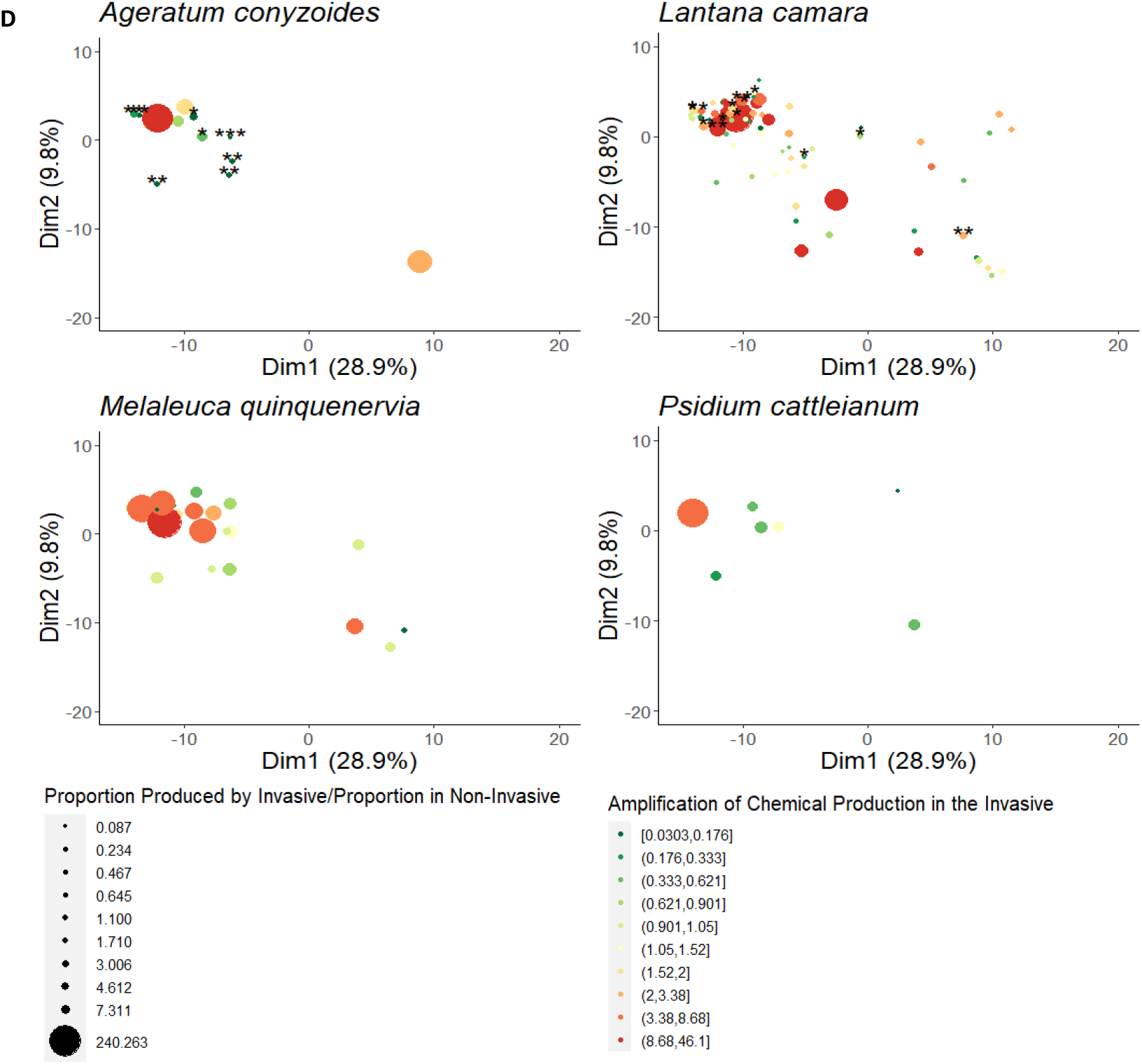
Differences in compound production between invasive compared to non-invasive populations, or non-native compared to native populations. Compounds are grouped according to K-medoids clustering from chemical distances. (a) Table indicating the overlap in clusters between the compounds in the species matrix compared to the family matrix. (b) Species table. The first part of the table shows the number of compounds shared between populations but with differing production levels. * ≤ 0.05 and ** ≤ 0.01 p-values are from exact binomial tests comparing the proportion of compounds represented by each cluster with different production between populations compared to the proportions of compounds belonging to each cluster produced in total by the invasive/non-native population. The second part of the table shows differences in unique-to-population compound production. * ≤ 0.05, ** ≤ 0.01, *** ≤ 0.001 and **** ≤ 0.0001 p-values from χ^2^ tests (left significance level on All Profiles) and exact binomial tests (right and for individual clusters) comparing compound synthesis between populations in total or per chemical cluster. Where two different significance levels are given, chi-square tests on unique-to-population compound synthesis for all clusters have also been computed. (c) Family table. The first part of the table shows the unique-to-population compound production with all profiles, as in (b). The second part of the table shows the average unique-to-population compounds produced and average p-values when the matrices were bootstrapped so that sample sizes from each population were equal. (d) Chemical distribution of compounds shared between invasive and non-invasive, or non-native and native, populations from the species matrix according to relative production levels and presence of the compounds in each form. * ≤ 0.05, ** ≤ 0.01 and *** ≤ 0.001 p-values from Mann-Whitney tests on compound production levels between the invasive and non-invasive form. The axes and compound coordinates were determined by K-medoids clustering using pairwise chemical distances between compounds using Euclidean distance methods. The x-axis explains 28.9% and the y-axis 9.8% of the variance seen in chemical distances data. Amplification of chemical production in the invasive was calculated from comparing production rates in invasives and non-invasives that did produce the compound.

**Fig 4.**
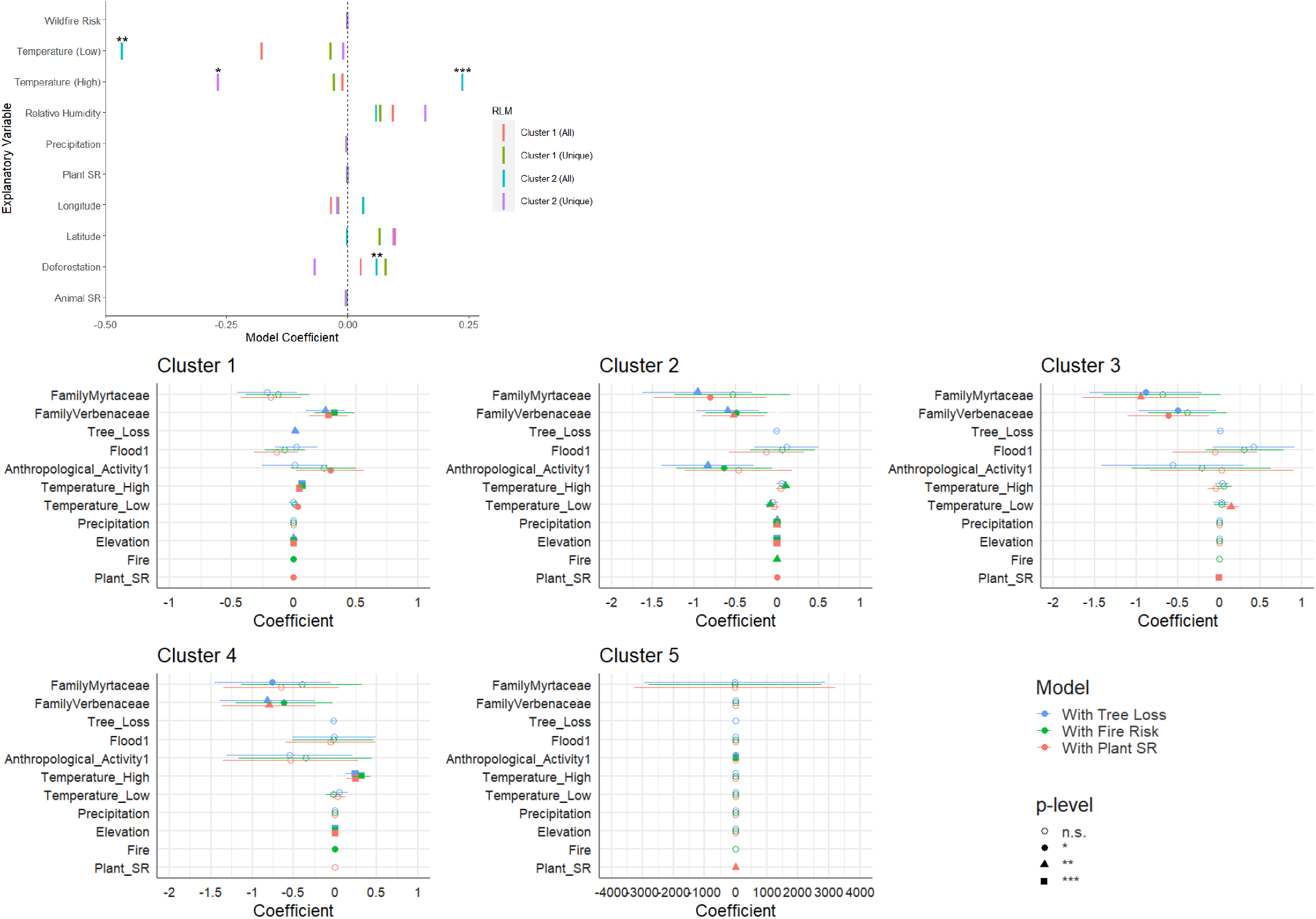
Compound distribution and production levels from invasive/non-native populations according to environmental and ecological factors. Coefficients from Robust Linear model (RLM) of average production rate of cluster 1 and 2 compounds per *L. camara* profile and average production of unique-to-invasive compounds from cluster 1 and 2 by environmental and ecological factors. * ≤ 0.05, ** ≤ 0.01, *** ≤ 0.001 p-values from Ward tests on how successfully explanatory variables modelled production. (See Materials and Methods for details on the explanatory variables.) (b) Coefficients from Poisson Regression evaluating the enrichment of clusters 1-5 against plant families and environmental and ecological factors. Three separate models were made so co-linear variables were not put in the same model nor removed from the data.

High numbers of compounds unique to populations, so were found exclusively in one population only, were found in non-native/invasive and native/non-invasive populations across all taxa (Fig. 3b, 3c). Trends were inconsistent across taxa; *L. camara* (*Verbenaceae*), *A. conyzoides* and *P. cattleainum* had significantly more compounds unique to invasive/non-native populations than non-invasive/native populations, whereas *M. quinquenervia, Myrtaceae* and *Asteraceae* showed the opposite. Patterns shown for all compounds were generally consistent across chemical clusters, although cluster 3 and 5 from the family matrix were always enriched in invasive/non-native populations. However, when equal sample sizes from the family data were compared using bootstrapping methods, generally the average number of unique-to-population compounds were greater in invasive than non-invasive populations, except for *Asteraceae*. Where compounds were enriched in invasive populations, cluster 2, 3, 5 and 6 were enriched, whereas only cluster 1 was enriched in taxa where native populations had more unique compounds. This was evidence for the average p-value from multiple exact tests. Therefore, chemodiversity radiates or contracts in invasive/non-native populations, but low-bias analysis methods suggest radiation is more common and invasive and native populations show radiation in different chemical structures.

### 3.5 Total variation in invasive secondary metabolomes does not relate to non-native environments and ecology, but metabolome chemical properties do

To understand some of the divergence observed in invasive populations, invasive profiles were compared to non-native environmental and ecological variables (Fig. 4). When comparing average compound production level per cluster in *L. camara* with environmental and ecological factors (Fig. 4a), cluster 1 production had little association with any factor. This was also shown using a robust linear model (RLM) and Ward tests. However, cluster 2 production associated non-significantly with *Plantae* and *Animalia* SR and significantly with deforestation rate, and average monthly high and low temperature. The same patterns were observed when only compounds unique-to-invasive-population, hence involved in novel invasive population evolution, populations were considered. Cluster 1 production associated very weakly with average low temperature and deforestation but was not significantly predicted by any explanatory variables. Cluster 2 production appeared to associate with temperature, fire risk, deforestation and *Plantae* and *Animalia* SR but was only significantly predicted by average monthly high temperatures. GLM regression analysis of cluster enrichment per profile across families also showed some compound production in invasive populations associated with environmental and ecological factors (Fig. 4b). Clusters 1, 2 and 4 associated with a variety of environmental and ecological factors across models, whereas clusters 3 and 5 rarely associated with any. Therefore, some variation in compound production and enrichment in structurally-similar compounds has evolved or responds plastically to non-native environments and ecology, so such compounds could be involved in local adaptation.

## 4. Discussion

The study of invasive plant evolution is well established, but scientific consensus on how these plants flourish in non-native environments is lacking (Zheng *et al*., 2015). For all hypotheses of how this occurs, including the EICA and NWH, secondary metabolomes are critical in driving emergence of invasivity, as secondary metabolites mediate most plant-environment interactions. In the emergence of invasivity, plant interactions with other plants, herbivores and other organisms are lost, gained and improved to increase fitness. However, studies into plant invasivity are constrained by not investigating complete secondary metabolomes, removing compounds of interest and additive effects and interactions between metabolites. We investigated the complete variation of secondary metabolomes, or chemical profiles, of invasive and non-invasive, or native and non-native, plants in one of the first attempts to correlated chemotype to phenotype. Evolution from native to invasive phenotypes was measured by analysing changes in production, loss and gain of metabolites between non-invasive to invasive and native to non-native populations. We demonstrate studying plant chemotype reveal novel insights on plant invasivity evolution, opening directions for new research.

### 4.1 There is high variation between all secondary metabolomes, but broad evolutionary strategies are conserved in invasives

Variation between the chemical profiles from all taxa was huge in terms of chemical composition (Fig. 1). Consistent evolutionary divergence between invasive and non-invasive populations and convergence within populations was expected, which should be reflected in chemical profile composition, but large variation was observed within as well as between populations. Additionally, total chemodiversity was biased by unequal sample sizes, where populations with greater sample sizes almost always had the greater chemodiversity. Despite this variation and confounding factors, consistent, but general, changes in secondary metabolome evolution can be observed in some taxa. Invasive individuals from *L. camara* and *Asteraceae* had undergone an expansion in chemodiversity, potentially diversifying plant-environment interactions. *L. camara* and *Achillea millefolium*, one of the non-native species in the *Asteraceae* family dataset, and has allelopathic and toxic properties (Rayaihi *et al*., 2002; Sousa *et al*., 2011); if the new compounds are involved in these processes, then NWH is proven. *M. quinquenervia* and *Myrtaceae* plants instead lost chemodiversity, suggesting specialist defence compounds have been lost and physiology and development has been invested in, so EICA could be true. Although such investment has been observed (Mishra, 2015), these results are unexpected as non-invasive populations have the greatest chemodiversity of any species, so further metabolic radiation is expected. Perhaps most of this chemodiversity can be attributed to specialist defence or metabolite genes were lost stochastically from small founder populations due to drift. *A. conyzoides* and *P. cattleainum* did not show any chemodiversity changes, so metabolites may have been lost and gained to equal degree, which could occur if NWH and EICA acted together. Therefore, metabolic evolution has occurred in invasive populations and EICA and NWH could be true to varying degrees based on the species.

### 4.2 Conserved changes in the functional properties of metabolomes could induce invasivity in accordance with EICA and NWH

Although conserved changes in the composition of chemical profiles were not observed (Fig. 1, Fig. 2), we hypothesised changes in compound production and the metabolome’s chemical properties to be a stronger signal for invasivity. This is because EICA states specialist defence compounds should be downregulated and the NWH states allelopathic compounds should be upregulated. Additionally, since it appears there is some stochasticity in which compounds undergo change in the invasive, there should be some functional convergence in chemotype, which could occur through structurally similar compounds being targeted for evolution. Cluster analysis showed compound production levels and chemical similarity of profiles were stronger signals of invasivity than chemical composition alone. (Fig. 2). Although there was not strong segregation of invasive and non-invasive populations, invasive and non-invasive phenotypes tended to group more separately into sub-clusters, suggesting some divergence had occurred. Within sub-clusters, non-invasive populations showed greater intra-population similarity than invasive populations, implying convergent evolution might not be important to invasives, instead the radiation of metabolomic diversity was. However, in *L. camara*, profiles clustered similarly when using production or chemical similarity data, suggesting there may be functional convergence in invasive plant metabolomes. This is further shown in other species when individual compound production was analysed individually and clustered according to chemical similarity. There was an overrepresentation of compounds with altered production in cluster 1 (from the species data), lost in *M. quinquenervia*, significantly downregulated *A. conyzoides* and *L. camara* and significantly upregulated in *L. camara* invasive populations (Fig. 3). Since EICA and NWH suggest compounds of similar function, defence and allelopathy, should be down- and upregulated respectively, the evidence suggests cluster 1 compounds are involved in these functions. This is further evidenced by significantly upregulated cluster 1 compounds in *L. camara* that have been bio-assayed – β-pinene and 1,8-cineole – are involved in allelopathy (Mishra, 2015). Moreover, multiple chemical families have multi-kingdom effects, involved in defence and allelopathy (Hickman *et al*., 2021) - cluster 1 compounds could have such effects. Therefore, structure-function relationships of metabolites have shown invasive metabolomes show evolutionary functional convergence despite diversifying. The EICA is also probably true for most species and NWH for *L. camara*.

### 4.3 Radiation in invasive chemodiversity increases fitness of the initial invasive phenotype and facilitates local adaptation

There was still unexplained variation in invasive metabolomes, so we hypothesised this was caused by radiation in metabolites involved in allelopathy, as predicted by NWH **(Torchin et al. 2003)**. Analysis of the full and bootstrapped data suggested most taxa had more unique metabolites in invasive compared to non-invasive populations. Many unique compounds clustered with the compounds with altered production in cluster 1, thought to be involved in allelopathy and defence, in the invasive populations all species. Contrastingly, chemical clustering from the family data suggested the alternative cluster 1, which has strong overlap with cluster 1 from the species data (Fig. 3a), was not enriched and instead depleted in invasive and enriched in non-invasive populations. However, alternative cluster 2 and 3 were enriched in invasive populations and had overlap with cluster 1 from the species data. Alternative cluster 2 and 3 also overlap with species cluster 2, which does contain allelopathic compounds like β- and γ-curcumene (Kato-Noguchi and Kurniadie, 2021). These combined results suggest some structurally-related compounds are involved in allelopathy and defence and allelopathy that is specialist or generalist. in invasive populations, there could have been loss of some specialist interactions and evolution of unique metabolites derived from retained more generalist metabolites. This implies the NWH could be true for many taxa.

However, the expansion of other clusters is not fully explained by changes in allelopathy or generalist defence strategies. To explain this expansion, and other unexplained variation between invasive profiles, evolution of invasives in response to environmental and ecological factors was also hypothesised. Modelling average production of clusters of all and unique-to-invasive compounds did show environmental and ecological factors influence invasive chemical profiles (Fig. 4a). Cluster 2 production was weakly impacted by environmental and ecological factors, but cluster 1 was not. The same pattern was observed for unique compounds. This implies some local adaptation was occurring and cluster 2 was not involved directly in general invasive aggressive strategies. It further proves cluster 1 compounds were involved in allelopathy or other aggressive invasive behaviour because evolution of these metabolites would be independent of the environment and ecology if all invasive plants utilise these strategies and all non-native species are susceptible (Callaway and Ridenour, 2004). However, modelling cluster enrichment within invasive families showed most clusters correlated with environmental and ecological factors (Fig. 4b). However, alternative cluster 3, which was enriched in invasive populations, was not correlated with environmental factors, suggesting these compounds were involved in generalist invasive strategies as predicted. Alternative clusters 1 and 2 may still be involved in more specialist interactions or have more diverse roles. Therefore, invasive plants do adapt to local non-native environments, so the emergence of invasivity is not just driven by aggressive strategies that are independent of the environment.

Therefore, an evolutionary framework can be built (Fig. 5). (1) After a plant is released from the selection pressure of specialist herbivory in a non-native environment, defence compounds used against these herbivores are downregulated and lost. (2) This reduction in metabolic and genetic load allows the invasive to invest in optimising pre-existing strategies to increase competitive advantage. If the plant already has a diverse metabolome, it could increase production of compounds involved in generalist herbivore defence, allelopathy, or other plant-environment interactions. Alternatively, the invasive may further lose chemodiversity, either neutrally or beneficially, if metabolites were specialised to its native range or if the plant had low initial chemodiversity initially and invest in physiological and developmental processes that directly increase growth and reproduction. (3) Increased competitive advantage increased resource acquisition. If the invasive invested in secondary metabolism, secondary metabolites will evolve and radiate from the pre-existing invested-in pathways. This is an environment-independent strategy. In parallel, secondary metabolism evolves in response to the local environment. Environment-independent and -dependent strategies combine to produce a high fitness phenotype. This theory needs, however, needs validation by testing other species.

**Fig 5.**
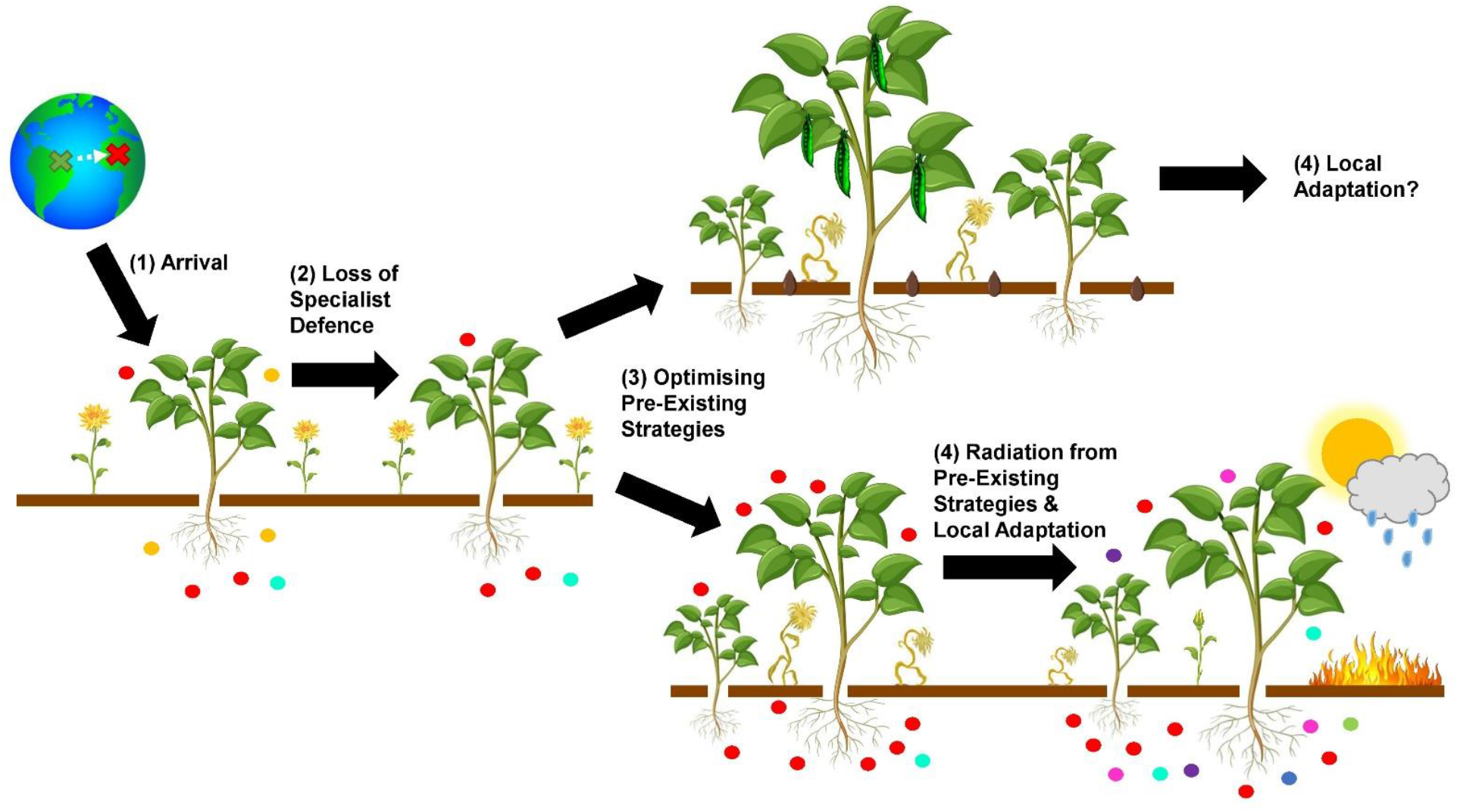
Schematic depicting plastic responses in the phenotype and the evolutionary trajectory of invasive plants.

## 5. Conclusions and Future Directions

There are many theories on how invasivity emerges in plants. We discovered, using computational methods and -omics approaches, secondary metabolomes of invasive plants have diverged from non-invasive populations and each other when native selection pressures are removed. Although diversification may be important in invasive evolution, invasive metabolomes appear to functionally converge to optimise invasivity potential, demonstrating a chemotype-to-phenotype relationship. Whether this was in accordance with the EICA, NWH or both varied between taxa, showing invasive evolution occurs across a spectrum of current evolutionary hypotheses. There was also evidence of local adaption to non-native environments. However, very few of the metabolites from this study have been assayed, which should be done in future to confirm these insights. Additionally, because of the lack of large datasets per taxa many computational methods like supervised machine learning could not be used, which would have been used to remove noise from unimportant compounds and identify specific compound sets driving invasivity. With such methods, potential for invasivity in non-native plants could be identified from metabolomes alone before the phenotype is apparent, a useful tool for conservationists. Therefore, improved data collection of essential oils and environmental surveying is needed so robust computational methods can be used to solidify the conclusions made in this study. Overall, however, this study emphasises the scope of multi-omics and computational approaches in producing novel insights into invasive plant evolution.

## Acknowledgements

The authors would like the thank the Plant Sciences Department, University of Cambridge for facilitating the creation of this project and for providing support.

## Author Contributions

J.R.C and G.Y conceived and designed this study. J.R.C, E.S and S.B collected and extracted the data. J.R.C did the computational and statistical analysis. J.R.C wrote the manuscript.

## Data Availability Statement

All data and code is available at https://github.com/jamilasrc/Quantifying-the-Invasive-Metabolome

## References

1. Blossey, B., & Notzold, R. (1995). Evolution of increased competitive ability in invasive nonindigenous plants: A hypothesis. The Journal of Ecology, 83, 887. https://doi.org/10.2307/2261425

2. Callaway, R. M., & Ridenour, W. M. (2004). Novel weapons: invasive success and the evolution of increased competitive ability. Frontiers in Ecology and the Environment, 2, 436–443. https://doi.org/10.1890/1540-9295(2004)002[0436:NWISAT]2.0.CO;2

3. Cappuccino, N., & Arnason, J. T. (2006). Novel chemistry of invasive exotic plants. Biology Letters, 2, 189–193. https://doi.org/10.1098/rsbl.2005.0433

4. Castro, M. R., Victoria, F. N., Oliveira, D. H., Jacob, R. G., Savegnago, L., & Alves, D. (2015). Essential oil of Psidium cattleianum leaves: antioxidant and antifungal activity. Pharmaceutical Biology, 53, 242–250. https://doi.org/10.3109/13880209.2014.914231

5. Copyright Holiday Weather-A-Connect Ltd. (n.d.). World Weather Averages. Weather2visit.Com; Weather 2 Visit. Retrieved December 3, 2021, from https://www.weather2visit.com

6. GBIF. (n.d.). Gbif.Org. Retrieved December 3, 2021, from https://www.gbif.org

7. Hickman, D. T., Rasmussen, A., Ritz, K., Birkett, M. A., & Neve, P. (2021). Review: Allelochemicals as multi-kingdom plant defence compounds: towards an integrated approach. Pest Management Science, 77, 1121–1131. https://doi.org/10.1002/ps.6076

8. Hull-Sanders, H. M., Clare, R., Johnson, R. H., & Meyer, G. A. (2007). Evaluation of the evolution of increased competitive ability (EICA) hypothesis: loss of defense against generalist but not specialist herbivores. Journal of Chemical Ecology, 33, 781–799. https://www.doi.org/10.1007/s10886-007-9252-y

9. Inderjit Callaway, R.M., & Vivanco, J. M. (2006). Can plant biochemistry contribute to understanding of invasion ecology? Trends in Plant Science, 11, 574–580. https://doi.org/10.1016/j.tplants.2006.10.004

10. Invasive Species Compendium. (n.d.).Cabi.Org. Retrieved December 3, 2021, from https://www.cabi.org/ISC

11. Kato-Noguchi, H., & Kurniadie, D. (2021). Allelopathy of Lantana camara as an invasive plant. Plants, 10, 1028. https://doi.org/10.3390/plants10051028

12. Kouame, B. K. F. P., Toure, D., Kablan, L., Bedi, G., Tea, I., Robins, R., Chalchat, J. C., & Tonzibo, F. (2018). Chemical constituents and antibacterial activity of essential oils from flowers and stems of ageratum conyzoides from ivory coast. Records of Natural Products, 12, 160–168. http://doi.org/10.25135/rnp.22.17.06.040

13. Kumari, S., Pundhir, S., Priya, P., Jeena, G., Punetha, A., Chawla, K., Firdos Jafaree, Z., Mondal, S., & Yadav, G. (2014). EssOilDB: a database of essential oils reflecting terpene composition and variability in the plant kingdom. Database: The Journal of Biological Databases and Curation, 2014, bau120. https://doi.org/10.1093/database/bau120

14. Kurade, N. P., Jaitak, V., Kaul, V. K., & Sharma, O. P. (2010). Chemical composition and antibacterial activity of essential oils of Lantana camara, Ageratum houstonianum and Eupatorium adenophorum. Pharmaceutical Biology, 48, 539–544. http://doi.org/10.3109/13880200903193336

15. Lind, E. M., & Parker, J. D. (2010). Novel weapons testing: are invasive plants more chemically defended than native plants? PloS One, 5, e10429. https://doi.org/10.1371/journal.pone.0010429

16. Mitchell, C. E., & Power, A. G. (2003). Release of invasive plants from fungal and viral pathogens. Nature, 421, 625–627. https://doi.org/10.1038/nature01317

17. Mishra, A. (2015). Allelopathic properties of Lantana camara. International Research Journal of Basic and Clinical Studies, 3, 13–28. http://dx.doi.org/10.14303/irjbcs.2014.048

18. Monti, D., Tampucci, S., Chetoni, P., Burgalassi, S., Bertoli, A., & Pistelli, L. (2009). Niaouli oils from different sources: analysis and influence on cutaneous permeation of estradiol in vitro. Drug Delivery, 16, 237–242. https://doi.org/10.1080/10717540902896297

19. Parker, J. D., & Hay, M. E. (2005). Biotic resistance to plant invasions? Native herbivores prefer non-native plants. Ecology Letters, 8, 959–967. https://doi.org/10.1111/j.1461-0248.2005.00799.x

20. Philippe, J., Goeb, P., Suvarnalatha, G., Sankar, R., & Suresh, S. (2002). Chemical composition of Melaleuca quinquenervia (cav.) S.T. Blake leaf oil from India. Journal of Essential Oil Research, 14, 181–182. https://doi.org/10.1080/10412905.2002.9699817

21. Qin, R.-M., Zheng, Y.-L., Valiente-Banuet, A., Callaway, R. M., Barclay, G. F., Pereyra, C. S., & Feng, Y.-L. (2013). The evolution of increased competitive ability, innate competitive advantages, and novel biochemical weapons act in concert for a tropical invader. The New Phytologist, 197, 979–988. https://doi.org/10.1111/nph.12071

22. Rayamajhi, MB., Van, TK., Center, TD., Goolsby, JA., Pratt, PD., Racelis, A. (2002). Biological attributes of the canopy-held Melaleuca seeds in Australia and Florida, US. Journal of Aquatic Plant Management, 40, 87–91.

23. Riaz, M., Khalid, M. R., & Chaudhary, F. M. (1995). Essential oil composition of Pakistani Ageratum conyzoides L. Journal of Essential Oil Research, 7, 551–553. https://doi.org/10.1080/10412905.1995.9698584

24. Senator, S. A., & Rozenberg, A. G. (2017). Assessment of economic and environmental impact of invasive plant species. Biology Bulletin Reviews, 7, 273–278. https://doi.org/10.1134/S2079086417040089

25. Siemann, E., & Rogers, W. E. (2003). Herbivory, disease, recruitment limitation, and success of alien and native tree species. Ecology, 84, 1489–1505. https://doi.org/10.1890/0012-9658(2003)084[1489:HDRLAS]2.0.CO;2

26. Sousa, S. M., & Viccini, L. F. (2011). Cytotoxic and genotoxic activity of Achillea millefolium L., Asteraceae, aqueous extracts. Revista Brasileira de Farmacognosia: Orgao Oficial Da Sociedade Brasileira de Farmacognosia, 21, 98–104. https://doi.org/10.1590/S0102-695X2011005000022

27. Strayer, D. L., Eviner, V. T., Jeschke, J. M., & Pace, M. L. (2006). Understanding the long-term effects of species invasions. Trends in Ecology & Evolution, 21, 645–651. https://www.doi.org/10.1016/j.tree.2006.07.007

28. Tesch, N. R., Mora, F., Rojas, L., DÍaz, T., Velasco, J., Yánez, C., Rios, N., Carmona, J., & Pasquale, S. (2011). Chemical composition and antibacterial activity of the essential oil of Lantana camara var. moritziana. Natural Product Communications, 6, 1031–1034. https://doi.org/10.1177/1934578X1100600727

29. The weather year round anywhere on earth. (n.d.). Weatherspark.Com. Retrieved December 3, 2021, from https://weatherspark.com

30. Torchin, M. E., Lafferty, K. D., Dobson, A. P., McKenzie, V. J., & Kuris, A. M. (2003). Introduced species and their missing parasites. Nature, 421, 628–630. https://doi.org/10.1038/nature01346

31. Vizzuality. (n.d.). Forest monitoring, land use & deforestation trends. Globalforestwatch.Org. Retrieved December 3, 2021, from https://www.globalforestwatch.org

32. weatherandclimate.com. (n.d.). Weatherandclimate.Com. Retrieved December 3, 2021, from https://www.weatherandclimate.com

33. Zheng, Y.-L., Feng, Y.-L., Zhang, L.-K., Callaway, R. M., Valiente-Banuet, A., Luo, D.-Q., Liao, Z.-Y., Lei, Y.-B., Barclay, G. F., & Silva-Pereyra, C. (2015). Integrating novel chemical weapons and evolutionarily increased competitive ability in success of a tropical invader. The New Phytologist, 205, 1350–1359. https://doi.org/10.1111/nph.13135

34. Zoghbi, M. das G. B., Bastos, M. de N. do C., Jardim, M. A. G., & Trigo, J. R. (2007). Volatiles of inflorescences, leaves, stems and roots of Ageratum conyzoides L. Growing wild in the north of Brazil. Journal of Essential Oil-Bearing Plants, 10, 297–303. https://doi.org/10.1080/0972060X.2007.10643558

